# Placozoan microbiomes reveal evolutionary trajectories towards mutualism among the largely parasitic *Rickettsiales*

**DOI:** 10.1101/2025.02.27.640636

**Authors:** Anna Mankowski, Henry Berndt, Nikolaus Leisch, Hiroaki Nakano, Genichiro Nishi, Michael G. Hadfield, Bruno Hüttel, Tina Enders, Nicole Dubilier, Harald R. Gruber-Vodicka

## Abstract

*Rickettsiales* are an enigmatic clade of strictly host-associated bacteria with some very common phylotypes in aquatic habitats. Despite amplicon- and metagenomic sequencing-based observations, the roles of these aquatic *Rickettsiales* are still unknown, as their genomes feature both beneficial and parasitic traits. In placozoans, microscopy-based observations of intracellular and *Rickettsia*-like-organisms point to *Rickettsiales* as common and abundant symbionts. We therefore systematically studied the microbiomes of placozoans to understand the eco-evolutionary dynamics of their associated *Rickettsiales*. Single-animal metagenomics revealed a widespread association of placozoans with *Rickettsiales* from three clades of ‘*Ca*. Midichloriaceae’, which were present in all but two placozoan lineages. A comparative phylogenetic analysis of placozoans and their *Rickettsiales* symbionts revealed host species-specific switches among *Rickettsiales* clades in several placozoan lineages. Our combination of evolutionary analyses, genome annotations, and expression data indicate different functional roles of these *Rickettsiales* clades, with some clearly diverging from a classical parasitic lifestyle by the loss of all ATP scavenging machinery. One of placozoan lineages that lacked ‘*Ca*. Midichloriaceae’ was instead associated with *Rickettsiaceae* symbionts, while another was completely devoid of *Rickettsiales* symbionts. Our imaging data of fibre cells entirely free of *Rickettsiales* underscores previously unrecognized variation in placozoan–microbe associations. Early-branching invertebrates, including placozoans as well as many corals, are gaining increasing traction as emerging models across the life sciences. Our data and analyses show that it is important to consider the specific endosymbiont groups and their ecology in a given host, as these symbionts likely influence the hosts’ cellular and organismal biology.

## Introduction

Several clades of bacteria are fully adapted to a host-dependent lifestyle, with all known members characterized exclusively as symbionts of other organisms. One of these enigmatic clades is the order *Rickettsiales*, alphaproteobacterial symbionts that occur intracellularly across a wide range of eukaryotic hosts, including many animals. They are generally regarded as parasites and cause a variety of diseases in humans, livestock and wild animals (*1, 2*). Although *Rickettsiales* are well characterized from terrestrial systems and insect hosts, their associations with marine invertebrates and protists have received comparatively less attention.

A key host group of marine *Rickettsiales* are placozoans, a phylum of globally distributed, benthic invertebrates that emerged very early in animal phylogeny (*3–5*). With their simple, two-layered bauplan and limited obvious morphological differentiation, placozoans were long considered a monotypic ‘phylum-of-one’. Molecular taxonomic studies first using mitochondrial 16S rRNA genes (haplotypes) and later comparative genomic analyses of placozoan genomes, alongside diversity patterns in other early-branching metazoans such as Cnidaria and sponges, have recognized four placozoan orders and six families. These include four formally described genera (*Trichoplax, Hoilungia, Polyplacotoma*, and *Cladtertia*) and three putative genera not yet formally described. The analyses of mitochondrial haplotypes indicated a large diversity of cryptic species (haplotypes) that are morphologically indistinguishable within placozoan genera (*3, 6–8*). Microscopy studies have consistently revealed that all placozoans examined harbor intracellular bacteria inside the fiber cells endoplasmic reticulum (*6, 9–11*). This widespread association between placozoans and intracellular bacteria, initially described from microscopy data, was also corroborated with genomic data in two placozoan lineages, and together with correlative light- and electron microscopy data indicates that these symbionts are *Rickettsiales* (*6, 9, 11–14*). This genomic data and symbiont specific imaging data for associated *Rickettsiales* so far came from two of the most closely related placozoan lineages from the genus *Trichoplax* (*14, 15*). In *Trichoplax adhaerens* (H1) and *Trichoplax* sp. ‘H2’, two closely related *Rickettsiales* genera from the family ‘Ca. Midichloriaceae’, ‘*Ca*. Grellia’ and ‘*Ca*. Aquarickettsia’ (*Midichloriaceae, Grellia* and *Aquarickettsia* from here on) were identified (*6, 13, 14*). However, the nature of the association between these two *Midichloriaceae* symbiont lineages and their placozoan hosts remains elusive, as their genomes appear to contain both mutualistic and parasitic traits (*14, 16*).

In this study, we analyzed the microbiomes of placozoans with an emphasis on putatively intracellular symbionts and specifically *Rickettsiales* by systematically sampling 115 single-animal metagenomes across the phylum Placozoa from nine sampling sites, representing three of the four orders of placozoans (*3, 17*). We combined a reconstruction of the host phylogeny with a systematic profiling of the microbiome of each individual, using primer-free and metagenome-based reconstruction of full-length host mitochondrial and bacterial 16S rRNA genes. We then focused on the *Rickettsiales* symbionts, analyzed their phylogenetic relations as well as their distribution across host haplotypes and performed last common ancestor reconstructions to recapitulate the evolutionary history between placozoans and *Rickettsiales* symbionts. We found an unexpected diversity with four different clades of *Rickettsiales* symbionts linked to host haplotypes, with host population specific association patterns across sites.

## Results & Discussion

### Placozoans are associated with a large diversity of putative symbionts

To describe the microbiome of placozoans across a broad biogeographic host range, we used single animal DNA extractions to generate ultra-low input shotgun DNA libraries from 115 host individuals from eight host mitochondrial 16S rRNA genotypes (host haplotypes) sampled in nine globally distributed locations (Fig. 1A-D, Table S1). We identified host haplotypes by reconstructing full-length mitochondrial 16S rRNA gene sequences, the standard in placozoan molecular taxonomy, and by inferring phylogenetic relationships between our samples and reference sequences (Figure 1D, Figure 2) (*3*). To characterize host-associated microbiomes, we first generated a database of *de novo* assembled 16S rRNA sequence from all metagenomes. Reconstructed sequences were summarized at the family level, except for *Rickettsiales*, which were split into genus-level taxa due to our specific research focus (Table S2). This database was then used for reference-based sequence reconstruction and to estimate the relative abundance of each taxon in individual hosts. Potential chimeras, likely human-or flow-cell–derived contaminants, and singleton families were removed, and relative abundances were normalized for downstream analyses (Note S1).

**Figure 1.**
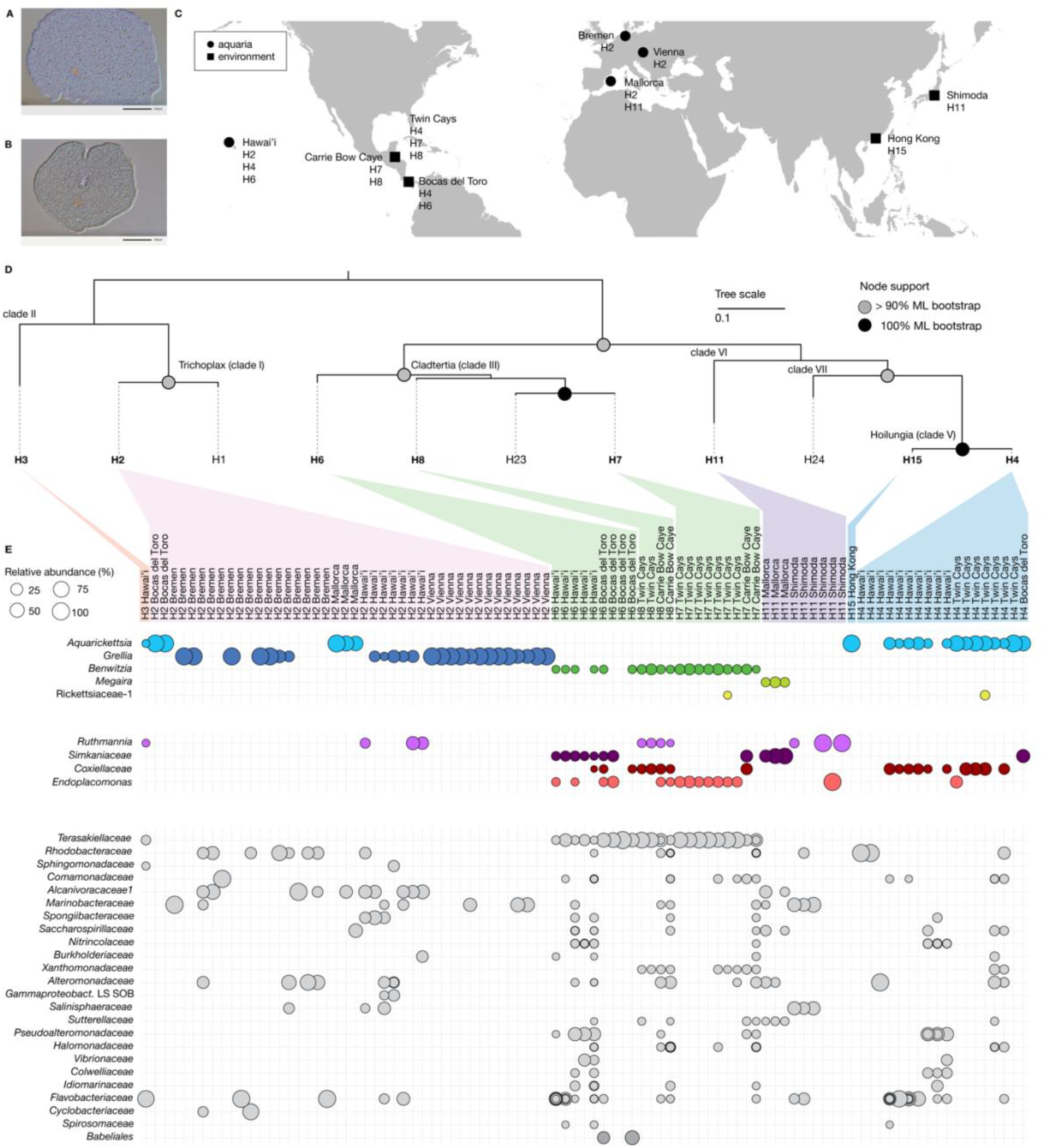
Placozoan individuals are associated with diverse microbiomes. A: DIC micrographs of a *Trichoplax* sp. H2 sampled in Vienna, Austria. B: DIC micrograph of a placozoan H11 sampled in Shimoda, Japan. C: World map showing sampling sites and detected host haplotypes. Circles indicate samples from aquaria, squares samples from the natural environment. D: Maximum-likelihood trees of the host mitochondrial 16S rRNA gene using one individual per host haplotype, with haplotypes analyzed in this study highlighted in bold. The scale bar indicates 10% estimated sequence divergence. Nodes with bootstrap support > 90% are shown in grey and 100% support in black. E: Relative abundances of all associated bacteria per host individual, as estimated with EMIRGE based on full-length 16S rRNA sequences from metagenomes. Each column represents a single host individual, and each row lists the identified bacterial group to the left, and shows abundances as dot plots scaled by relative abundance. Rows are arranged to show intracellular taxa on top, starting with members of the *Midichloriaceae*. Note that almost all host individuals have detectable amounts of *Rickettsiales* endosymbionts, largely *Midichloriaceae*. Most other bacterial families are irregularly distributed, and are often restricted to specific host clades, e.g. the *Endoplacomonas. Rickettsiales* symbionts are colored by genus; other putative endosymbionts are colored by class; all other detected bacteria are shown in gray.

**Figure 2.**
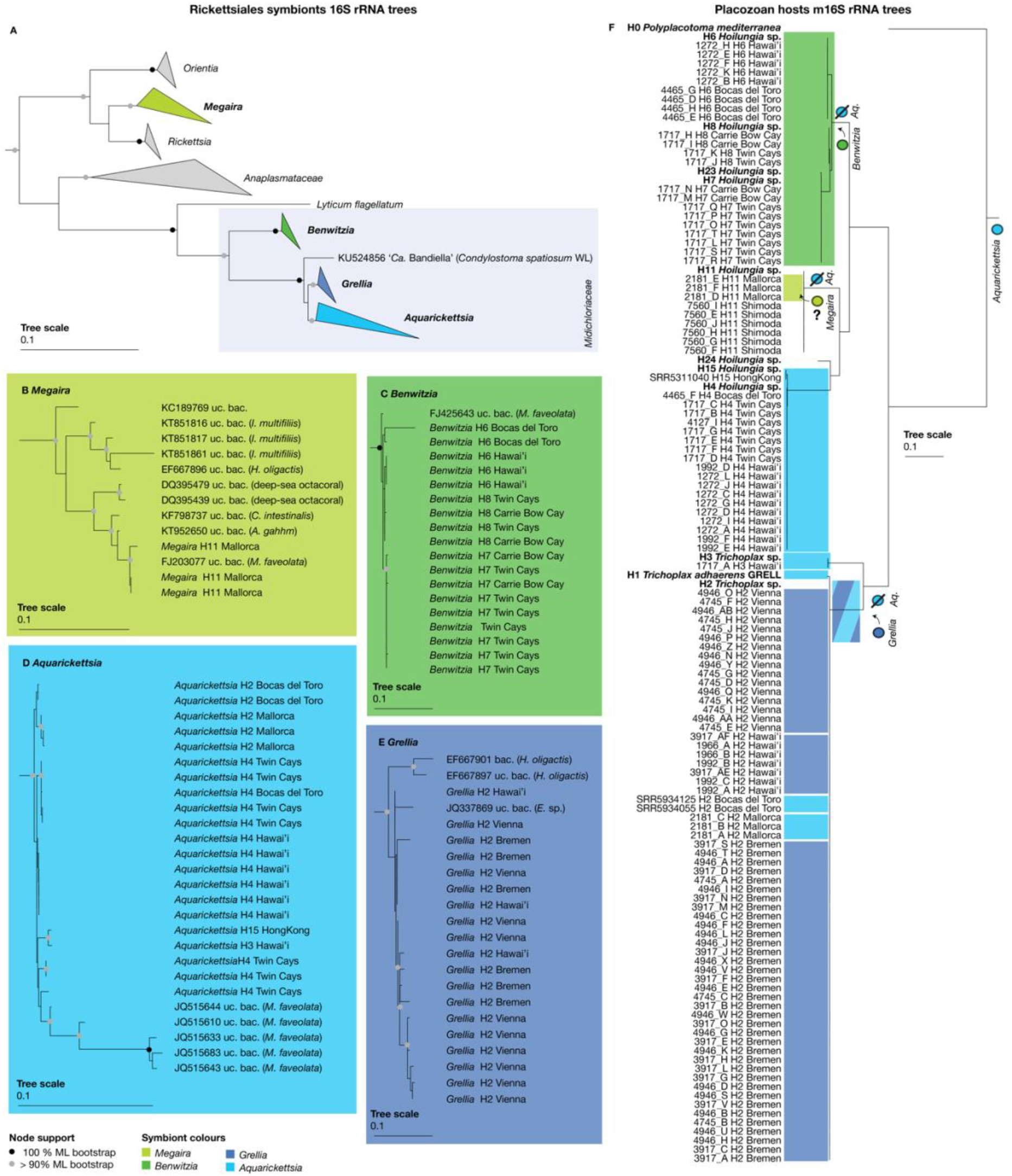
Phylogeny of placozoan associated *Rickettsiales* (A-E) and their distribution and evolutionary history across the host diversity (F). A-E: 16S rRNA based phylogenetic reconstructions. A-F: scale bars indicate 10% estimated sequence divergence A: overview of the *Rickettsiales* symbionts and close relatives. Colored triangles highlight the different *Rickettsiales* symbiont genera. B: *Megaira* symbionts and close relatives. C: *Benwitzia* symbionts and close relatives. D: *Aquarickettsia* symbionts and close relatives. E: *Grellia* symbionts and close relatives. F: m16S rRNA Phylogeny of the host individuals. References are highlighted in bold. Colored boxes highlight the predicted association with the different *Rickettsiales* symbiont clades based on a maximum-parsimony model. Predicted events of symbiont losses and acquisitions are indicated at the corresponding nodes of the host phylogeny. Small cartoons illustrate the inferred symbiont lineages associated with each split based on the last common ancestor reconstruction.

Our microbiome screening revealed a widespread and consistent association of placozoans with *Rickettsiales* symbionts (Fig. 1), including phylotypes that belong to the previously described *Grellia* and *Aquarickettsia* clades (*13, 14, 18*). The screening of many different haplotypes and sampling sites however revealed an even larger diversity of placozoan-associated *Rickettsiales*. In total, we detected five genera specifically associated with different haplotypes from different sampling locations (Fig. 1, Fig. 2). In addition, we detected intracellular bacteria from the *Margulisbacteria*, ‘*Ca*. Ruthmannia’ (*Ruthmannia* from here on) in three placozoan haplotypes from four locations (Fig. S1D). *Ruthmannia* symbionts have been previously characterized with genomic, expression and microscopy data in the H2 haplotype (*14*). Their association with placozoans appears patchy but more common than anticipated (*19*), as they were repeatedly present across haplotypes and sampling sites. We additionally detected three more families of putative endosymbionts which have not been described as placozoan symbionts yet. These include two clades of *Simkaniaceae* (Fig. S1A) and one clade of *Coxiellaceae* (Fig. S1B), both known to occur solely as intracellular symbionts of diverse eukaryotes (*20, 21*). In addition, we detected one clade of *Endozoicomonadaceae* (Fig. S1C), a family of common symbionts of marine invertebrates (*22–24*). The *Endozoicomonadaceae* detected in placozoans are only distantly related to the well-characterized genus *Endozoicomonas* and represent a new genus we name ‘*Ca*. Endoplacomonas’ (*Endoplacomonas* from here). While there is a high chance that these three families are yet uncharacterized endosymbionts of placozoans, their position in the animals and their roles still needs genomic reconstructions and microscopy-based detection, e.g., via fluorescence *in situ* hybridization using taxon-specific probes.

Beyond the detection of five families of putative endosymbionts, we detected 24 additional families. 23 of these 24 families had a broad geographic or taxonomic range of hosts, but occurred only in a few individuals, which might indicate a non-specific association, sequencing bias or contamination, e.g., members of the *Flavobacteriaceae*. In contrast, the *Teraskiellaceae (Alphaproteobacteria*) were only detected in certain haplotypes (H3, H6, H7 and H8), but consistently across all specimens, which could indicate a specific association. However, further investigation and additional data are necessary to understand the nature of their relationships to placozoans.

Our microbiome characterization revealed a so far undescribed diversity of bacteria associated with placozoans, in varying degrees of specificity across host haplotypes and sampling sites. For many of them, the type of association as well as their role and impact on placozoans remains unclear, but, given their phylogenetic background, five of the detected families are very likely intracellular endosymbionts. While most bacterial representatives were associated with a few individuals/haplotypes, the association with *Rickettsiales* symbionts is almost omnipresent across the diversity of placozoan hosts. We only detected a single lineage (H11 from Shimoda, Japan) in which none of the specimens was associated with *Rickettsiales* symbionts (Figure 1, Table S3). In all other lineages, we detected *Rickettsiales* in at least a subset of the specimens. For many specimens in which we could not assemble a full-length 16S rRNA gene, we nevertheless detected medium to high proportions of reads mapping to *Rickettsiales* reference genomes (Table S3). This indicates that *Rickettsiales* were likely present but could not be assembled due to low sequencing coverage. In addition to *Rickettsiales*, no more than two other putative endosymbiotic taxa were detected per individual. It remains to be shown if they are cell-type specific and do not co-occur, as was shown for *Grellia* and *Ruthmannia* in H2 (*14*), or if they overlap in certain cell types in some host-symbiont combinations.

### Diversity, distribution and evolutionary history of *Rickettsiales* symbionts

Given the widespread and consistent association between placozoans and *Rickettsiales*, we next explored the hypothesis that the *Rickettsiales* are linked to patterns of placozoan evolution. To this end we excluded the genus ‘Rickettsiaceae-1’ from our downstream analyses, as this phylotype was detected only sporadically and in relatively low abundance, and always co-occurred with a higher abundant and more broadly distributed *Rickettsiales* symbiont, in contrast to the widespread and high abundance association between the placozoans and all other *Rickettsiales* phylotypes. For the remaining and widespread assocations, our microbiome screening revealed four *Rickettsiales* genera. This includes the previously described *Aquarickettsia* and *Grellia* symbionts. Both taxa belong to the *Midichloriaceae* and are the sister clade to ‘*Ca*. Bandiella’. In addition, we also detected two more genera of *Rickettsiales* symbionts that have not been described in placozoans yet. The first one, we name ‘*Ca*. Benwitzia’ (*Benwitzia* from here) and also belongs to the *Midichloriaceae*, while the fourth one is a *Rickettsiaceae* from the genus ‘*Ca*. Megaira’ (*Megaira* from here). All four genera have been reported from other marine protists and animals, indicating a broad distribution of *Rickettsiales* across diverse marine hosts (Fig. 4)(*25–27*). Consistent with this, in our phylogenetic analyses, the *Rickettsiales* symbionts detected in placozoans are closely related or even form intermixed clades with symbionts of other marine invertebrates, most often corals, for which the association with *Aquarickettsia* has been studied extensively (Figure 2)(*13, 18*).

We detected a *Rickettsiales* phylotype in all haplotypes from all sampling sites, except one lineage of H11 from Shimoda, Japan, and could reconstruct a full-length 16S rRNA for nearly all individuals. This widespread association between the different Rickettsiales clusters was highly specific for certain haplotypes and dominated by *Midichloriaceae*, such as *Aquarickettsia* that were most widespread and associated to most haplotypes (H1, H2, H3, H4 and H15). Members of *Grellia* were much rarer and were only associated with lineages of the H2 haplotype from Bremen, Vienna and Hawai’i, while members of the *Benwitzia* clade were found across the monophyletic clade of H6, H7 and H8 haplotypes. Only a single host haplotype (H11) did not host *Midichloriaceae*, but was either associated with *Megaira* symbionts from the *Rickettsiaceae* (lineage from Mallorca) or did not feature any *Rickettsiales* symbionts at all (isolates from Shimoda).

Our analyses indicate that *Rickettsiales* associations are ancestral in placozoans but highly dynamic, with multiple independent replacement and loss events across lineages. *Aquarickettsia* appears to be the primary symbiont, retained across many recent haplotypes, reflecting the ancestral state of the association of placozoans with *Rickettsiales* symbionts (Table S3 and S4). In some lineages, however, this symbiont has been replaced. In the *Cladtertia* clade (haplotypes H6, H7, and H8), the consistent presence of *Benwitzia* across all sampled populations suggests that this symbiont replaced *Aquarickettsia* in the common ancestor of the clade, and that this replacement was subsequently inherited by descendant haplotypes. Similarly, in haplotype H2, three lineages (Bremen, Hawai’i, and Vienna) show replacement of *Aquarickettsia* by *Grellia*, whereas populations from Bocas del Toro and Mallorca retain *Aquarickettsia*. Other cases are less clear. In haplotype H11, *Aquarickettsia* is absent in all specimens from Mallorca and Shimoda, and *Megaira* is present in the Mallorca population. We cannot determine whether the loss of *Aquarickettsia* occurred once in a common ancestor or independently in each population, nor whether the acquisition of *Megaira* preceded or followed this loss. Likewise, because the host phylogeny based on mitochondrial 16S rRNA does not fully resolve relationships among lineages, it is unclear whether the replacements observed in H2 populations represent independent events or were inherited from a shared ancestor. Overall, these patterns indicate that placozoan-*Rickettsiales* associations involve both long-term retention of ancestral symbionts and repeated host-switching or replacement events, reflecting a dynamic evolutionary history that remains incompletely resolved.

### H11 individuals that lack symbionts in the rough endoplasmic reticulum reveal placozoan origin of mitochondrial complexes in fiber cells

To corroborate the symbiotic status of the *Rickettsiales*-free H11 population from Shimoda, we performed differential interference contrast (DIC) microscopy on living and transmission electron microscopy (TEM) on high pressure frozen specimens (Fig. 3, Fig. S2-3). Generally, the microanatomy of H11 resembled the general placozoan body plan (*12, 28–30*). In concordance with the absence of any *Rickettsiales* symbionts from the metagenomic data, endosymbionts in the endoplasmic reticulum of fiber cells were not observed. (Fig. 3, Note S2). The consistent presence of *Rickettsiales* in the rER of fiber cells, together with the peculiar mitochondrial complexes observed in all placozoan species, led to the hypothesis that these intracellular symbionts might induce the unique mitochondrial morphologies that have not been reported in any other eukaryote. Here, we show that the formation of these mitochondrial stacks occurs independently of *Rickettsiales*, indicating that they are an intrinsic feature of the placozoan cellular bauplan (*12*).

**Figure 3.**
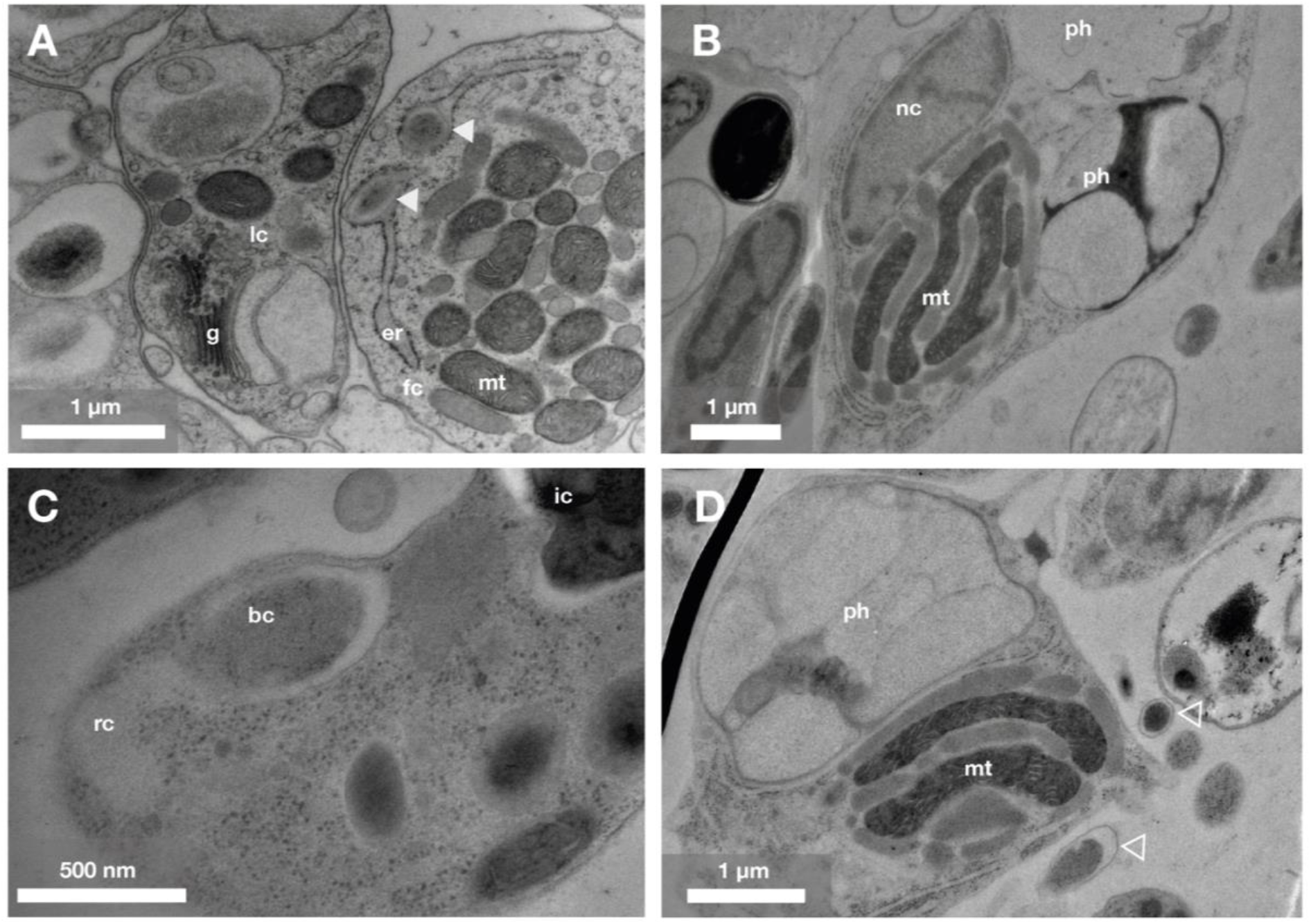
The placozoan H11 from Shimoda shows the typical mitochondrial complexes in its fiber cells, but this lineage lacks the endosymbionts assumed to induce this phenotype. A: Transmission electron microscopy (TEM) micrograph of a fiber cell of *Trichoplax* sp. H2 ‘Vienna’ containing *Grellia* in the endoplasmic reticulum (er, closed arrowheads). lc: lipophilic cell; g: golgi apparatus; mt: mitochondria. B:TEM micrograph of a fiber cell of the placozoan H11 ‘Shimoda’ without endosymbionts in the ER. nc: nucleus; mt: mitochondria; ph: phagosome. C: TEM micrograph of an intracellular bacteria (bc) in the cytoplasm of a cell of the lower epithelium of H11. rc: reorganized cytoplasm; ic: inclusion. D: Micrograph of a fiber cell of H11 with extracellular bacteria next to it (open arrowheads). ph: phagosome; mt: mitochondria.

**Figure 4.**
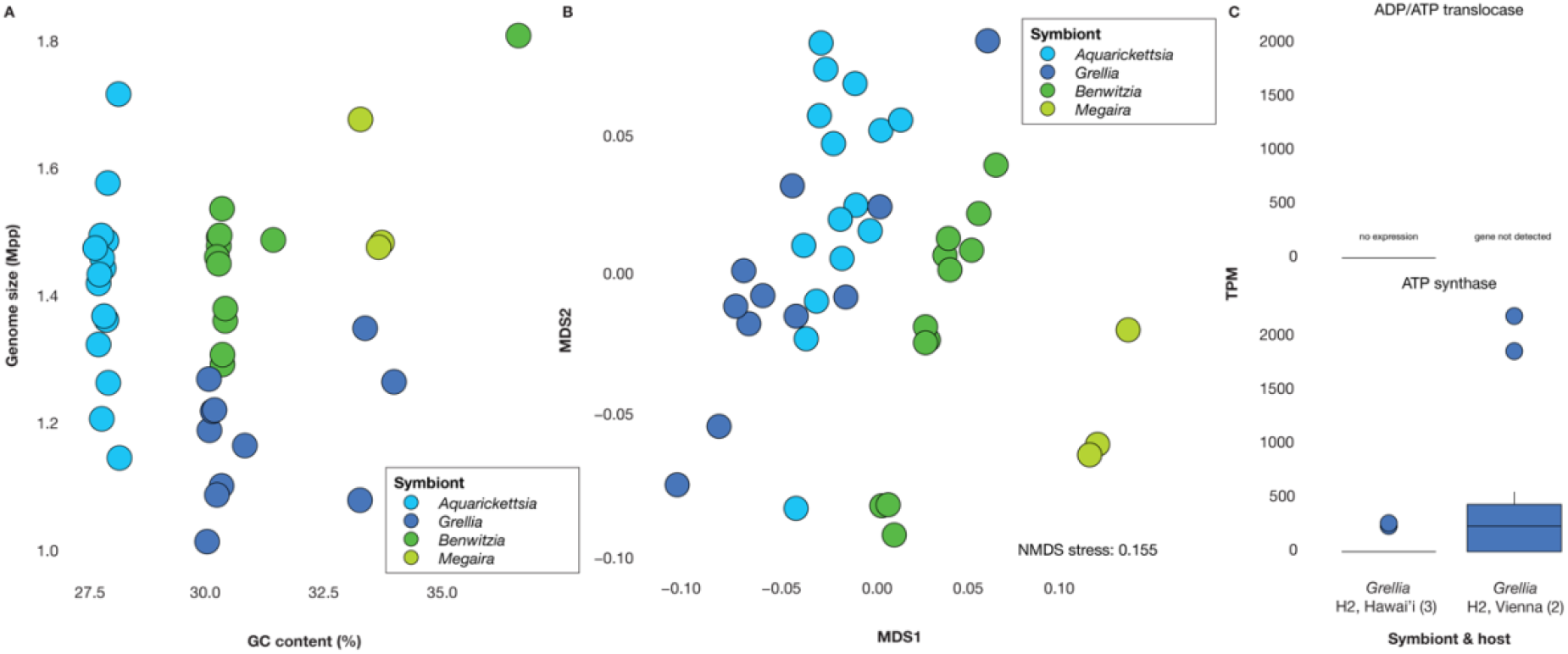
Genomic and transcriptomic analyses of *Rickettsiales* symbionts indicate diverging functional profiles and a convergent loss of energy parasitism. A: Genome size (Mbp) and GC content of *Rickettsiales* genomes. Points are colored by symbiont type. B: NMDS plot of Bray-Curtis distances calculated based on the frequency of COG categories in *Rickettsiales* symbiont genomes. Points are colored by symbiont type. C: Transcription of genes for ADP/ATP translocases and ATP synthases estimated through pseudoalignment of transcriptomic reads to reference MAGs. Boxes are colored by symbiont type. Numbers in parentheses refer to the number of analyzed metatranscriptomes.

Despite the absence of *Rickettsiales* symbionts in this population, we observed intracellular bacteria in cells of the lower epithelium (Fig. 3C, Fig. S3J-L) and extracellular bacteria in the intra-epithelial region of H11 (Fig. 3D, Fig. S3H). The identity of these bacteria remains unknown and they may be food-related, loosely associated bacteria or true symbionts. Metagenomic analyses indicate that *Ruthmannia* are the only abundant symbionts in all specimens from this population and morphologically, the extracellular bacteria resemble the previously characterized *Ruthmannia* symbionts (*14*). This is consistent with observations of *Ruthmannia* in the ventral epithelium of other Placozoa haplotypes, however, in the case of H11, they seem to occur extracellularly in contrast to their intracellular localization in H2 specimen (*14*). Our morphological characterization shows that H11 from Shimoda displays the classic placozoan body plan, including distinctive mitochondrial stacks in fiber cells that appear to form independent of the presence of *Rickettsiales* symbionts.

### Genomic and transcriptomic analyses of *Rickettsiales* symbionts reveal an evolutionary trajectory to minimize the energetic burden for the host

The observation of host specificity in *Rickettsiales* symbionts led us to investigate their genomic features more closely. We therefore used metagenomic assembly and binning to generate metagenome assembled genomes (MAGs) of *Rickettsiales* symbionts from 42 samples with sufficient coverage.

Symbiont genomes with sufficient quality ranged from 1.0 to 1.96 Mbp, with GC contents between 28% and 37% (for MAGs with >50% checkM2 completeness, and <10% contamination, Table S4). While these genomic traits do not indicate the extreme genome reduction seen in many insect symbionts, they nonetheless reflect a host-associated lifestyle. There was no clear trend suggesting different levels of host dependence (i.e., smaller genome size and lower GC content). Instead, genomes of *Aquarickettsia* symbionts had the lowest GC content, whereas *Grellia* genomes were the smallest. Given their close phylogenetic relationship, these subtle but stable differences in genome evolution are intriguing and points towards differences in their evolutionary trajectories. In contrast, *Megaira* genomes were the largest and had the highest GC content, with *Benwitzia* symbionts showing intermediate genome statistics.

To assess whether functional capacities differ between these symbionts, we compared the number of COG (Clusters of Orthologous Genes) categories in each genome(*31*). Despite slight differences in genome size and GC content, *Aquarickettsia* and *Grellia* were most similar among the symbiont groups based on the frequency of different COG categories and did not form distinct clusters, indicating comparable functional fingerprints. Consistent with the trends in genome size and GC content, *Megaira* was the most distinct, with *Benwitzia* falling in between (PERMANOVA: R^2^ = 0.51, p = 0.001, ANOSIM: R = 0.70, p = 0.001). Although analyses of symbiont composition and evolutionary history (Figures 1 and 2) indicate that one symbiont may replace another, the observed differences in functional profiles show that these replacements do not result in complete functional equivalence.

To investigate the potential energetic burden of *Rickettsiales* symbionts on placozoan hosts, we examined their capacity for ATP parasitism. *Rickettsiales* are generally thought to scavenge ATP from their hosts via ADP/ATP translocases, while many can also generate ATP using their own ATP synthases. A genomic survey of placozoan-associated *Rickettsiales* revealed that all symbionts possess genes encoding ATP synthases, and most also carry ADP/ATP translocases (Extended Data). Notably, some *Grellia* lineages from H2 haplotypes sampled in Vienna and Bremen lack the translocase, indicating potential lineage-specific differences in ATP acquisition strategies (Extended Data, Figure S4).

To further assess functional activity, we compared expression of ATP synthase and ADP/ATP translocase genes in two *Grellia* lineages. Consistent with previous studies, only ATP synthase genes were detectably expressed, even when the translocase gene was present (*14*). These data suggest that, under the conditions examined, *Grellia* generates ATP primarily through its own synthase rather than the classical host-dependent ATP import, although occasional host-derived ATP import cannot be excluded. In contrast, *Megaira* symbionts from the H11 haplotype in Mallorca retain multiple ADP/ATP translocase variants, indicating that these symbionts rely more heavily on host ATP, consistent with a more classical parasitic strategy within the *Rickettsiaceae*. Overall, these findings highlight variation in ATP acquisition strategies among placozoan symbionts, with potential implications for the energetic burden imposed on the host, while emphasizing that functional conclusions remain tentative given the current data.

## Conclusion

This study provides a detailed inventory of the placozoan microbiome, revealing both previously characterized and novel symbionts. In addition to *Rickettsiales* and *Margulisbacteria*, we identify putative endosymbiotic associations with *Endozoicomonadaceae, Simkaniaceae*, and *Coxiellaceae*. We also catalog a range of microbes whose association with placozoans remains uncertain. These potentially loosely associated microorganisms offer opportunities to investigate their interactions and functional roles within the host.

Among these associations, *Rickettsiales* symbionts are abundant, widespread, and specific. Our reconstructions suggest that the association with a *Rickettsiales* partner is likely ancestral across modern placozoans. Phylogenetic analyses further indicate that placozoans may exchange symbionts with other marine invertebrates, as members of all four placozoan symbiont clades are intermixed with symbionts from Cnidaria, Porifera, and Ctenophora (Figure 2) (*13, 14*). A recent study predicted that *Aquarickettsia* symbionts of corals are horizontally transmitted, suggesting a free-living life stage with likely limited survival time(*14, 16, 18*)which could provide a source for acquisition in placozoans. Given these potential symbiont switches, and the observation that both *Aquarickettsia* and *Grellia* reside in the rER lumen (*9, 11, 12*), placozoans may serve as a model to study how intracellular, and even intra-organelle, symbioses are established.

Despite the widespread and potentially ancient association, it remains unclear whether placozoan Rickettsiales behave as classical parasites, as observed in reef-building corals, or whether their association is more commensalistic or mutualistic. Previous genomic and transcriptomic analyses show that *Grellia* and *Aquarickettsia* genomes harbor both mutualistic and parasitic traits (*13, 14, 16*). While Rickettsiales are often described as energy parasites reliant on host ATP(*32, 33*) our analyses indicate that *Grellia* and *Aquarickettsia* symbionts appear to generate ATP primarily via their own ATP synthases, with some lineages losing ADP/ATP translocases entirely. This pattern is rare among

*Rickettsiales*, previously observed only in lineages associated with protist hosts, where it has been proposed to reduce host dependence (*34*). An additional, non-exclusive possibility is that reliance on self-generated ATP minimizes energetic costs for the host. However, no comprehensive metabolic budgets are available, and the loss or absence of a translocase does not conclusively rule out occasional host ATP use. Notably, coral-associated *Aquarickettsia* often considered parasitic, also encode ATP synthases and may rely primarily on self-generated ATP, though expression has not been confirmed (*13*). Overall, these observations suggest that *Grellia* and *Aquarickettsia* diverge from classical *Rickettsiales* parasitism by reducing energetic dependence on their hosts, while the precise nature of these interactions remains unresolved. In addition, although both symbionts are auxotrophic for amino acids and nucleotides, they may supplement host diets with vitamins, such as riboflavin, which placozoans cannot synthesize (*14*). By contrast, *Benwitzia* and *Megaira* exhibit markedly different functional profiles, which may reflect more classical parasitic associations, including sustained ATP parasitism.

Placozoans have emerged as a promising model system for studying metazoan evolution, developmental biology, and tissue formation (*3, 35, 36*). Given their diverse microbiomes and the likely ancestral, widespread, and specific association with *Rickettsiales*, microbial partners should be considered in many research contexts. Most *Rickettsiales* are potent manipulators of host cellular and organismal biology, and the *Midichloriaceae*, with multiple secretion systems and numerous predicted effectors, could influence host development, for instance contributing to arrested early development during sexual reproduction (*10, 14*). At the same time, the intimate integration of intracellular symbionts with host organelles makes placozoans an ideal model for studying host-microbe interactions. Available single-cell atlases from several placozoan lineages (*37*) provide a framework to examine how intracellular *Rickettsiales* stabilize their niches, manipulate hosts, or contribute to host physiology.

## Material & methods

### Sample collection, processing and metagenomic and -transcriptomic sequencing

115 individual placozoans were sampled at various field sites (Table S1). Placozoans were identified using a dissection microscope and afterwards kept in culture in 400 ml glass beakers. For individuals kept in culture, the culture medium was 34.5‰ artificial seawater and cultures were fed weekly with 2×10^6^ cells ml^-1^ of *Isochrysis galbana*. Cultures were kept at 25°C and a 16:8 hours light:dark cycle. DNA was extracted from single individuals with the DNeasy Blood & Tissue Kit (Qiagen, Hilden, Germany) according to the manufacturer’s instructions with the following exceptions: the proteinase K digest was performed overnight, elution volumes were halved, all samples were eluted twice reusing the first eluate and all elutions were performed after ten minutes of waiting prior to centrifugation. Library construction, quality control and sequencing were performed at the Max Planck Genome Centre (Cologne, Germany). If needed, DNA concentration was increased using the MinElute PCR Purification kit (Qiagen). Libraries for Illumina sequencing were prepared using the Ovation Ultralow Library Systems Kit (NuGEN) according to the manufacturer’s instructions or using a miniaturized Tn5 transposase-base protocol including a limited PCR step (Table S1). The libraries were size selected by agarose gel electrophoresis. The quality and quantity of selected fragments were analysed by fluorometry and capillary electrophoresis on LabChip GXII Touch. Paired end reads of 100, 150 or 250 bp length were sequenced on an Illumina HiSeq 4000 or an Illumina NextSeq 2000 sequences (Table S1). In addition, we included three previously published placozoan metagenomes into our analysis. They were downloaded from the NCBI short read archive (accession numbers SRR5311040, SRR5934055 and SRR5934125).

Transcriptomes generated from *Trichoplax* sp. H2 from Hawai’i were obtained from ENA (project PRJEB30343). For each transcriptome from *Trichoplax* sp. H2 from Vienna, 20 individuals of a clonal *Trichoplax* sp. H2 strain isolated from a commercial aquarium in Vienna, Austria were processed using the Qiagen RNeasy Micro Kit following the manufacturer’s instructions for the isolation of RNA from animal tissue. RNA concentration was determined using the Thermo Fisher Qubit 4.0 with the Qubit RNA High Sensitivity kit. The purified RNA was stored at −80 °C until further processing for RNA sequencing. Samples were processed using the Illumina stranded total RNA kit with Ribozero Plus rRNA depletion (Illumina, San Diego, USA) according to the manufacturer’s manual. Sequencing was performed on the NovaSeq 6000 (Illumina, San Diego, USA) using 100bp paired-end sequencing.

### Assembly of host marker genes, phylogenetic inference and identification of host haplotypes

Mitochondrial 16S rRNA genes of all specimens were assembled by adapting the phyloFlash pipeline to operate on a custom m16S rRNA gene reference database and predict m16S rRNA genes from assembled sequences (*38*). The detected m16S gene sequences were aligned to m16S rRNA gene sequences of previously identified specimens using mafft-xinsi v7.407 (*3, 39–41*). Maximum-likelihood based phylogenies were calculated using IQ-TREE, including automatic selection of the best suited model and generation of 1000 ultrafast bootstrap replicates (*42*). The sequence of *Polyplacotoma mediterranea* H0 was used to root the phylogenies. Host haplotypes were defined based on the phylogenetic relations to the previously identified specimens (*3*).

### Symbiont clade definition and quantification

16S rRNA genes were assembled from the metagenomic libraries using phyloFlash using the – almosteverything option and in addition specifying the read length. All resulting bacterial sequences were inspected for chimera using the uchime algorithm implemented in vsearch v2.6.2 (*43*) and providing sequences matched in the first mapping step of the phyloFlash run as reference database. Only non-chimeric sequences were considered for downstream applications. The reference based on matched sequences from the phyloFlash run was additionally used to place phyloFlash assembled 16S rRNA gene sequences into a phylogenetic tree. The reference database was clustered at 95% sequence identity using the easy_clust algorithm from mmseqs2 (*44*). Assembled sequences and database hits were aligned using mafft-xinsi and a phylogenetic tree was calculated from the resulting alignment using IQ-TREE including automatic selection of the best suited model and generation of 1000 ultrafast bootstrap replicates. After verifying the phylogenetic placement, assembled sequences were grouped at the genus-level (for members of the Rickettsiales) or the family level (for all other taxa).

The abundances of all groups were quantified across all metagenomic libraries using EMIRGE v.0.61.1 following the standard workflow for custom EMIRGE databases (*45*). Chimeric sequences were identified as described above. Additionally, we confirmed phylogenetic placement for non-chimeric sequences by aligning them to the phyloFlash assembled sequences and the clustered collection of phyloFlash database hits using mafft-xinsi and calculating a phylogenetic tree with IQ-TREE (including automatic selection of the best suited model generation of 1000 ultrafast bootstrap replicates). Subsequently, we excluded chimeric sequences, taxa that appear to be contaminations or appeared only once (Note S1, Table S2). After contamination removal, we normalized the relative abundances of the remaining clades to 100%.

### Phylogeny of all symbionts and their relatives

Sequences of all putative symbiont clades were used to obtain sequences from closely related bacteria from the SILVA database (*46*). Here, we used the SINA search and classify algorithm to obtain up to 10 relatives for each sequence that shared at least 90% sequence similarity for each of our input sequences (*47*). For the Rickettsiales, we in addition screened the RefSeq database using BLAST implemented in Geneious v11.1.5 to obtain the ten most similar 16S rRNA genes (*48*). Duplicated sequences were removed from the collection of sequences of the symbionts’ relatives. The resulting sequence collection was aligned using mafft-xinsi and a phylogenetic tree was calculated using IQ-TREE including automatic selection of the best suited model generation of 1000 ultrafast bootstrap replicates.

### Last common ancestor analysis of the *Rickettsiales* symbionts

To determine the evolutionary history of the association of *Rickettsiales* symbionts with placozoans, we performed last common ancestor modeling with pastml (*49*). We used DOWNPASS prediction methods for a maximum-parsimony based estimate.

### Metagenomic binning and annotation

Symbiont MAGs were generated using a custom pipeline that combines trimming and filtering of raw reads, metagenomic assembly and binning. The exact pipeline is shared under: https://github.com/amankowski/MG-processing_from-reads-to-bins/tree/main. Bin quality was examined by genome completeness and contamination (checkM v1.2.2 (*50*)), the presence of a 16S rRNA gene (barrnap v0.9, https://github.com/tseemann/barrnap) and the number of amino acids that had at least one tRNA encoded in the genome as a second proxy for genome completeness (ARAGORN v1.2.38). Taxonomy was assigned to bins that were of at least medium quality according to MIMAG standards (> 50% complete and showed < 10% contamination, GTDBtk v1.3.0, GTDB r95 (*51, 52*)). If multiple bins were present per library and symbiont taxon, we selected the best one based on genome completeness and contamination estimates, the presence of 16S rRNA, and the number of amino acids with at least one tRNA encoded. We then removed contigs that were shared by at least two bins from the same library. Quality of the final bins of *Rickettsiales* symbionts was checked again with checkM v2 (*53*), barrnap and ARAGORN and the genome statistics of these final bins are reported in Table S4. Functional genome annotations were generated using bakta (*54*). The similarity of metabolism across *Rickettsiales* symbionts was inferred from clusters of orthologous groups (COG) categories, which represent broad functional classifications (*31*). We generated COG category profiles for each symbiont using eggNOG-mapper v2.1.6 (*55*) with DIAMOND alignment (*56*). To quantify functional similarity, we calculated the relative frequencies of COG categories and used them to compute Bray-Curtis distances between genomes. These distances were then visualized using non-metric multidimensional scaling (NMDS). Statistical differences among symbiont taxa were tested using PERMANOVA and ANOSIM with 999 permutations.

### *Rickettsiales* MAG coverage in sequencing libraries

As an additional measure to assess symbiont presence or absence, we mapped all metagenomic libraries against all MAGs generated in this study. We used coding sequences (CDS) predicted by bakta and read mapping was performed with kallisto v0.45.0 (*57*) with default settings against the full set of predicted CDSs. For each genome, we calculated the proportion of genes to which at least one read mapped, and for *Rickettsiales* genera represented by multiple MAGs, we report the mean proportion across all MAGs.

### Transcriptome analyses

We quantified transcription levels of ATP synthase and ADP/ATP translocase genes by mapping RNA-seq reads to CDSs of the highest quality *Rickettsiales* reference MAGs from the same host haplotype and sampling location from using kallisto v0.45.0 with default settings.

### Analyses and plotting of symbiont community composition

The analyses of symbiont community composition were performed in R v4.41 unless differently stated. During the analyses, the following packages were used: (https://github.com/sdray/ade4), tidyverse (*58*), ggplot2 from the tidyverse package, maps (https://www.rdocumentation.org/packages/maps), mapdata (https://www.rdocumentation.org/packages/mapdata).

### Light and electron microscopy

For light microscopy, placozoan individuals were mounted in artificial sea water (ASW) on glass slides and covered with coverslips for differential interference microscopy using a Leitz Orthoplan light microscope (Leitz, now Leica, Germany) equipped with 25X, 40X and 100X DIC objectives and a Gryphax NAOS 20 MP camera (Jenoptik, Germany). Overview images were recorded using the 25X objective (Air, NA 0.50) and high-magnification images were conducted using the 100X objective (Oil-Immersion, NA 1.30). For electron microscopy, animals were allowed to settle in a type A gold-coated copper sample carrier of 100 μm depth (Leica Microsystems, Germany). Subsequently, ASW was removed, the carrier was filled with 1-hexadecen, sealed with a type B gold-coated copper sample carrier and subjected to high-pressure freezing using a Leica EM ICE (Leica Microsystems, Germany). The frozen samples were placed on top of frozen acetone containing 0.1% osmium tetroxide in liquid nitrogen and freeze substituted (FS) using the Leica AFS2 freeze substitution unit. After 8 hours at −90°C, the temperature was ramped up to −60 °C within 8 hours, held for 8 hours at −60 °C and subsequently ramped up to −40 °C within 4 hours. The samples were incubated for one hour on ice in the FS cocktail, washed with acetone and were infiltrated in Spurr’s low viscosity epoxy resin using a fast embedding protocol (*59*).The blocks were cured at 60 °C for 48 hours. Sections were generated using diamond knives (Diatome, Switzerland) on a Leica EM UC7 ultramicrotome (Leica Microsystems, Germany). For light microscopy, 1 μm thin sections were generated that were collect on glass slides and stained with methylene blue. For transmission electron microscopy (TEM), 60 nm ultrathin sections were generated and collected on custom Formvar coated copper slot grids. Before TEM imaging, sections were contrasted 30 minutes with saturated aqueous uranyl acetate (UA) and 1 minute on Reynold’s lead citrate. TEM images were recorded using 3a FEI Tecnai G2 Spirit BioTWIN (FEI Company, now Thermo Fisher Scientific, USA) operated at 80 kV or 120 kV.

### Declaration of generative AI and AI-assisted technologies in the writing process

During the preparation of this work, the authors used ChatGPT (OpenAI) to improve language and clarity. After the usage of this tool, the authors reviewed and edited the contents as needed and take full responsibility for the content of the article.

## Author contributions

AM and HRGV conceived the study and AM analyzed data with input from HRGV and TE. HB generated imaging data of placozoan specimens. HRGV, NL, HN, GN, MGH, and BH acquired placozoan specimens and generated metagenomic data. AM wrote the manuscript together with HRGV and HB, with input from all co-authors.

## Availability of data and materials

Raw metagenomic and -transcriptomic sequences are deposited in the European Nucleotide Archive (ENA) under accession ID PRJEB107716 and will be made available upon publication of the manuscript and are currently available upon request. Host and symbiont marker genes as well as symbiont MAGs and annotations are deposited under https://doi.org/10.5281/zenodo.18410361 and will be made available upon publication. We deposited all phylogenetic trees as newick files through iTOL v6 (*60*). Data and scripts used for the generation of figures and statistical analyses are shared under https://github.com/amankowski/placozoan_microbiome.

## Acknowledgements

We are thankful for sample collections and field assistance by Chris Keller, Daniel Abed-Navandi, Emilia M. Sogin, Laetitia Wilkins, Ramon Rossello-Mora, Tomas Wilkop and Zach Foltz. In addition, we would like to thank the Carrie Bow Cay Laboratory and the Bocas del Toro Research Station of the Smithsonian Tropical Research Institute, the Haus des Meeres Vienna, the Las Palma Aquarium and their staff for supporting our sampling campaigns. We thank Sam Vohsen for constructive scientific discussions. We also thank the Central Microscopy (Kiel University) for their support with TEM acquisition (Urska Repnik). This work is contribution XXX from the Carrie Bow Cay Laboratory, Caribbean Coral Reef Ecosystem Program, National Museum of History, Washington DC.

