## Supplemental Figures and Notes, Extended Data for "Placozoan microbiomes reveal evolutionary trajectories towards mutualism among the largely parasitic *Rickettsiales*"

#### Supplementary tables (provided in a separate document)

Table S1: Overview over samples used in this study and additional metadata.
Table S2: Taxonomy of phyloFlash assembled 16S rRNA sequences.
Table S3: Overview of assembled 16S rRNA sequences and mapping proportions of reads to *Rickettsiales* genomes.
Table S4: Genome statistics of *Rickettsiales* MAGs.

#### Supplementary Notes

##### Note S1: Removal of chimeras, singletons and likely contaminants from the 16S rRNA sequence collection

To ensure the robustness and biological relevance of downstream phylogenetic and diversity analyses, we applied several filtering steps to remove artefactual and contaminant sequences. First, all assembled 16S rRNA gene sequences were screened for chimeras. Chimeric sequences can arise during assembly and represent artificial recombinations of multiple parental templates. If not removed, such sequences can inflate diversity estimates and introduce spurious phylogenetic signal. We screened for chimeric sequences both computationally (all EMIRGE assembled sequences) and by manually inspecting the phylogenetic tree (calculated with EMIRGE assembled sequences that were at least 1000 bp long and did not contain more than 20 ambiguous bases). During manual curation, we observed one phyloFlash-assembled sequence assigned to *Alteromonadaceae* that was placed within *Endozoicomonadaceae*. Because the alignment appeared sound, it was initially used as an EMIRGE reference. However, all sequences reconstructed from this reference clustered with *Alteromonadaceae*. We therefore concluded that the phyloFlash-assembled sequence was chimeric, and all EMIRGE-derived sequences based on it were discarded. In addition, we observed a few *Pseudomonadota* (*Gammaproteobacteria*) sequences appeared chimeric, as they did not cluster with reference sequences from their assigned family. Thus, these sequences were also flagged and removed from downstream analyses. Similarly, we observed some *Flavobacteriaceae* sequences that did not cluster with reference sequences from their assigned genus. These however were retained, as they remained within the correct family. Second, singleton taxa (i.e., taxa detected in only a single metagenomic sample) were excluded, as they are unlikely to represent biologically relevant or persistent symbionts and are more prone to reflect technical artefacts or contamination. Finally, sequences identified as human-derived or originating from sequencing flow cell contamination were removed. These contaminants are commonly introduced during library preparation or sequencing and do not reflect genuine biological associations of the host organism. Together, these filtering steps ensure that the retained dataset represents high-confidence microbial sequences and minimizes the impact of technical artefacts on downstream analyses.

**Note S2: Microanatomy and ultrastructural features of the placozoan haplotype 11 (H11) strain 'Shimoda'**

Based on differential interference contrast microscopy on live specimens of the placozoan H11 from Shimoda, it shares the same basic body plan and organization with other well characterized placozoans: a densely populated, ciliated lower epithelium (Fig. S2A-B), a dorsal epithelium consisting of wider, ciliated cells (Fig. S2C) and a region between the epithelia containing e.g. crystal cells (Fig. S2D-E). In the central region of the lower epithelium, cells with voluminous vesicles close to the apical membrane are apparent, which are likely lipohilic cells with lipophilic vesicles (lv, Fig. S2A)(*12*, *28*–*30*). The upper epithelium features multiple vesicles larger than lipophilic vesicles, which from their position and size are likely "shiny spheres" (SS) – structures derived from cells of the upper epithelium that are present in many placozoan haplotypes and apparently contain toxic effectors to repel predators (*61*). In the region between the epithelia, we observed crystal containing cells close to the periphery of the animal (Fig. S2D-E). These crystal cells (CC) have been shown to function as statocysts for gravity sensing (*12*, *62*). Additionally, we consistently observed dense accumulations of shiny-sphere-like vesicles to the outmost periphery of the animal (Fig. 2SD-E). Given the proposed function for anti-predator defense, this organization of shiny spheres at the outer borders of the animal could be an adaptation to predators seizing placozoans from the side close to the substrate, like e.g. slugs from the family Rhodopidae, which have been shown to feed on placozoans (*63*). Even more intriguingly was the observation of extracellular bacteria (bc) within the placozoan tissue in the inter-epithelial region (Fig. S2E).

As our metagenomic data on H11 specimens from Shimoda also indicated the absence of the typical intra-ER endosymbionts of the family *Midichloriaceae* but indicated the presence of other bacteria, we opted for ultrastructural analysis of the 'Shimoda' lineage. Using high-pressure freezing, freeze substitution and embedding in Spurr low viscosity epoxy resin, we generated thin and ultrathin sections for light and transmission electron microscopy (TEM, Fig. S3A-B). Also on the ultrastructural level, the resemblance with the typical placozoan body plan is evident (Fig. S3A-B). Dorsal epithelial cells (DEC and ventral epithelial cells (VEC) as main constituents of the lower and upper epithelium, as well as secretory lipophilic cells (LC) in the lower epithelium and fiber cells (FC) in the inter-epithelial region are easily distinguishable by their morphological habitus, i.e. the characteristic mitochondrial complex of fiber cells and the voluminous Golgi-derived vesicles of lipophilic cells (*12*, *29*). Focusing on cross-sectioned fiber cells to test the absence or presence of endosymbionts, we observed not a single intracellular bacterium in fiber cells of two different individuals of the placozoan H11 'Shimoda' (Fig. S3-D-G) confirming the observed absence of *Midichloriaceae* from metagenomic data. Instead, we observed extracellular bacteria within the intra-epithelial region (Fig. S3-H-I) first observed during DIC microscopy. These bacteria feature an inner and outer membrane, which indicates that they belong to a gram-negative bacterial clade. Additionally, we found intracellular bacteria within cells of the lower epithelium (Fig. S3-J-L). Close to the intracellular bacteria, we also observed host cytoplasm lacking ribosomes – an observation we preliminarily term 'reorganized cytoplasm'. The observed extracellular bacteria could either be a free-living state of these intracellular bacteria for e.g. colonization of new host cells or a distinct bacterial clade. To this end, the identity of the observed intra- and extracellular bacteria remains to be resolved, but the prevalence of *Ruthmannia* symbionts in the metagenomic data generated from H11 'Shimoda' specimen indicates that some of the observed bacteria should belong to this group. Morphologically, the extracellular bacteria seem to share more similarity with *Ruthmannia* symbionts from the H2 haplotype, however, these were described to be intracellular symbionts (*14*). Further experiments, such as 16S rRNA-targeted fluorescence *in situ* hybridization (FISH) with specific probes for different members of the microbiome of the placozoan H11 'Shimoda' could enable the identification of the associated bacteria and reveal whether the *Ruthmannia* symbionts occur intra- or extracellularly in H11 'Shimoda'.

### Supplementary figures

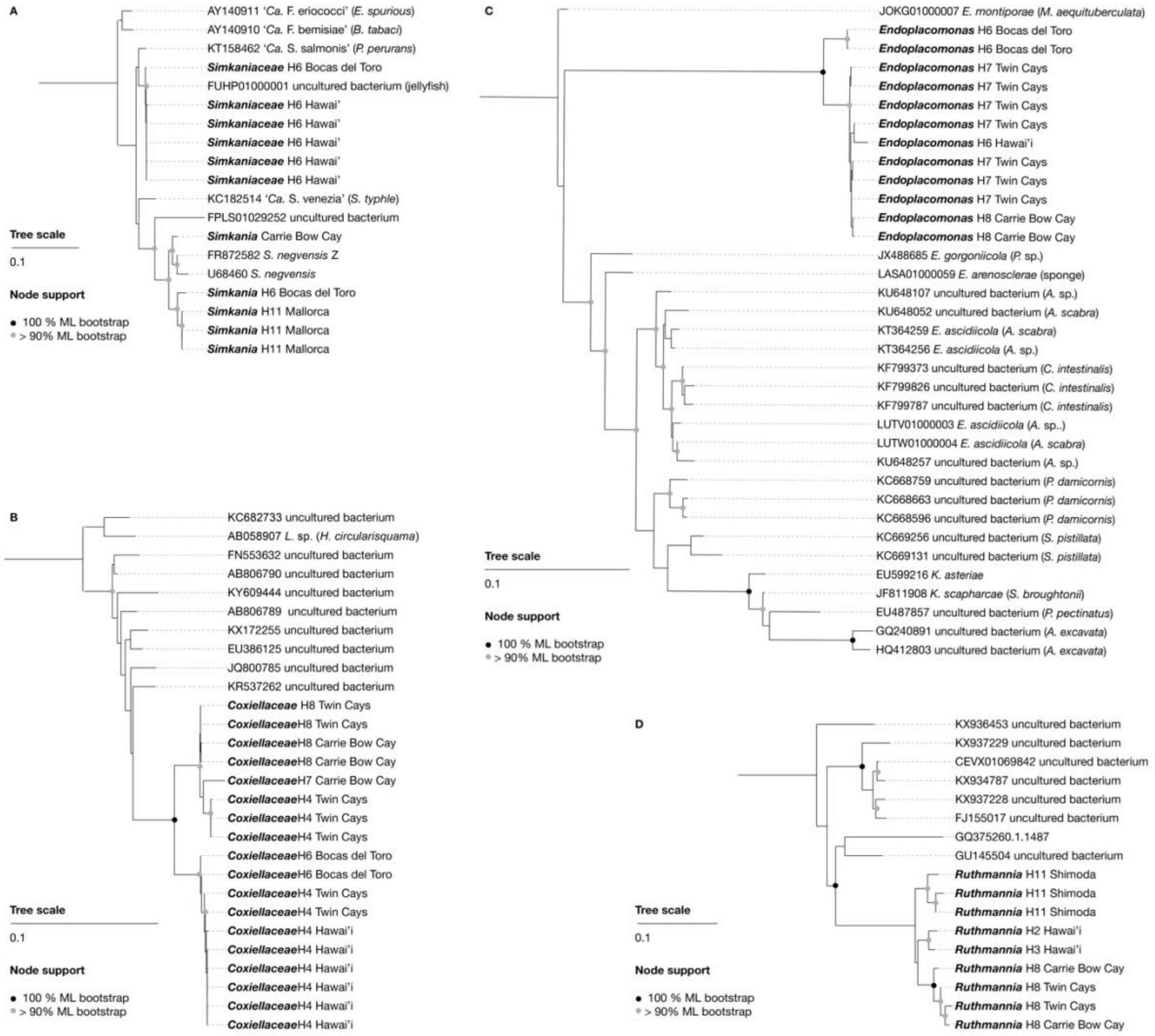


**Figure S1 | 16S rRNA phylogenies of putative endosymbionts of placozoans.** A: *Simkaniaceae* B: *Coxiellaceae* C: *Endozoicomonadaceae* D: *Margulisbacteria*

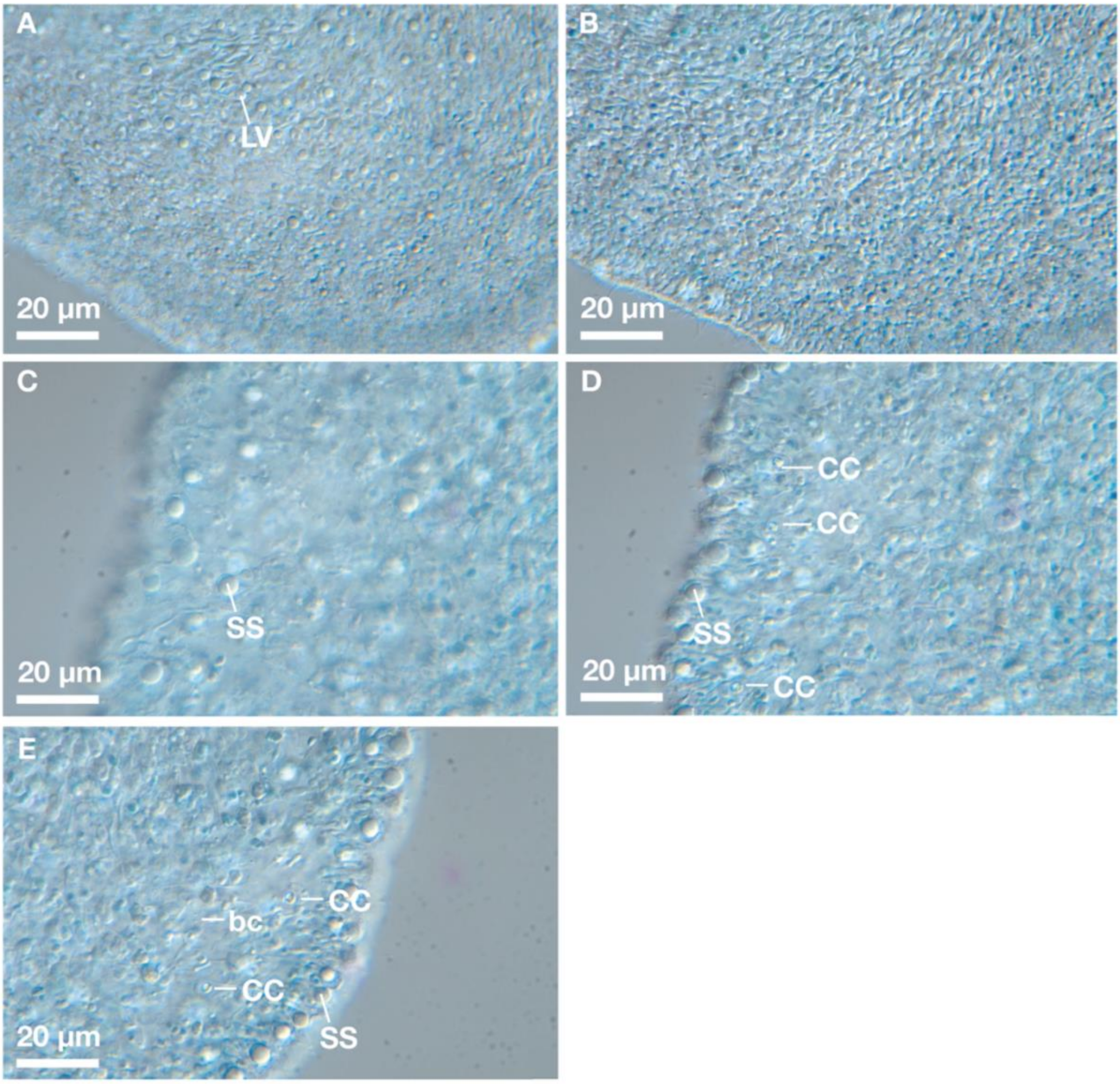


**Figure S2 | Microanatomy of the placozoan H11 'Shimoda' by DIC microscopy.** A: Micrograph of the apical regions of the lower epithelium featuring presumably lipophilic vesicles (LV) in lipophilic cells. B: Micrograph of the columnar cell bodies of the densely populated lower epithelium. C: Micrograph of the upper epithelial epithelium with multiple shiny spheres (SS). D: Micrograph if the inter-epithelial region with multiple crystal cells (CC) and a peripheral ring of shiny-sphere-like vesicles. E: Another micrograph of the interepithelial region that contains extracellular bacteria (bc).

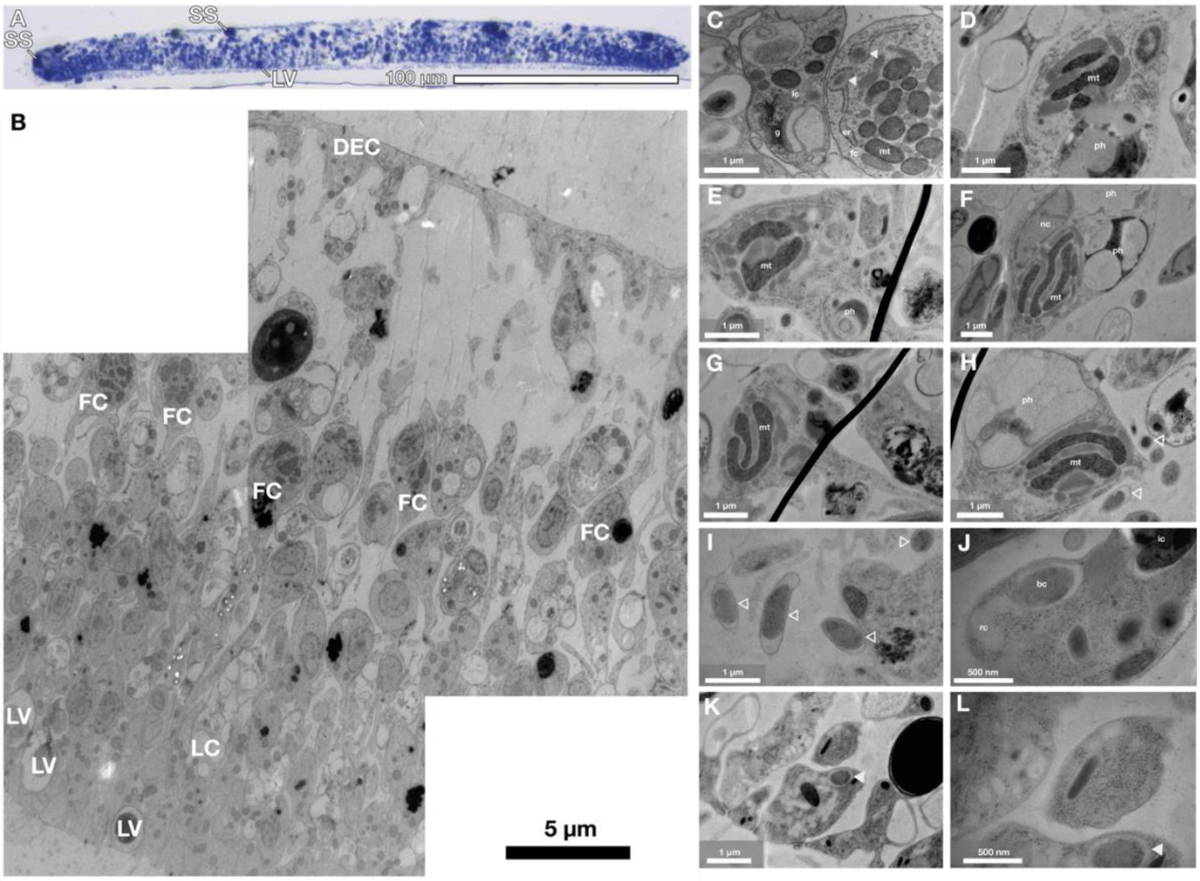


**Figure S3 | Microanatomy of the placozoan H11 'Shimoda' by TEM. A: 1 µm section stained with methylene blue.** Shiny spheres (SS) in the upper epithelium and at the periphery are distinguishable, as well as lipophilic vesicles (LV) in the lower epithelium. B: Manually stitched image from two TEM micrographs of H11. Dorsal epithelial cells (DEC), fiber cells (FC), lipophilic cells (LC) and lipophilic vesicles (LV) close to the apical membrane of lipophilic cells are annotated. C: Representative micrograph of a fiber cell of the placozoan *Trichoplax* H2 'Vienna', featuring *Grellia* in the endoplasmic reticulum (closed arrowheads). D-G: Representative micrographs of fiber cell from two H11 'Shimoda' individual lacking endosymbionts in the endoplasmic reticulum. H: Fiber cell of H11 'Shimoda' without endosymbionts but with extracellular bacteria next to it (open arrowheads). I: Micrograph of the observed extracellular bacteria in the inter-epithelial region (open arrowheads). J: Micrograph of a cell of the lower epithelium containing an intracellular bacterium (bc) with reorganized cytoplasm (rc) next to it. K: Micrograph of the lower epithelium with another cell containing an intracellular bacterium in the cytoplasm (closed arrowhead). L: Higher magnification of the cell shown in K. g: golgi apparatus; lc: lipophilic cell; fc: fiber cell; er: endoplasmic reticulum; mt: mitochondria; ph: phagosome; ic: inclusion; nc: nucleus; bc: bacterium; rc: reorganized cytoplasm.

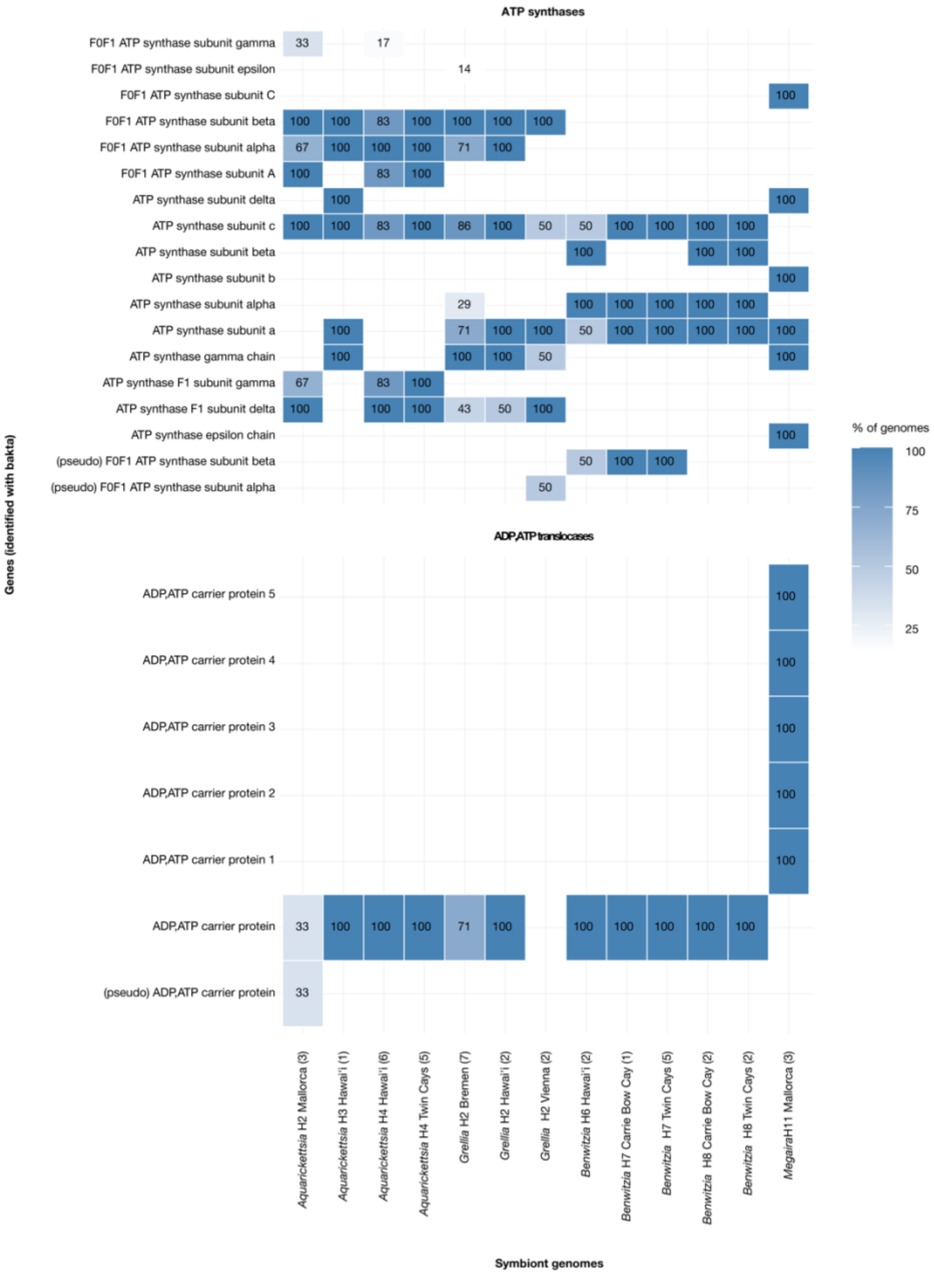


**Figure S4 | Percentages of MAGs from each placozoan haplotype and sampling location carrying different ADP/ATP translocase genes.** Genes were identified using bakta (*54*). Numbers in parentheses indicate the number of MAGs analyzed.

### Extended data

#### Phylogenetic trees

Phylogeny of host m16S rRNA genes from reference samples and all metagenomes from this study:
https://itol.embl.de/tree/10111897465771768490276

Phylogeny of all bacterial 16S rRNA genes reconstructed from metagenomes with phyloFlash as well as reference sequences:
https://itol.embl.de/tree/10111897192617684903 60

Phylogeny of all bacterial 16S rRNA genes reconstructed from metagenomes with EMIRGE as well as reference sequences:
https://itol.embl.de/tree/10111897331721768548752

#### rRNA sequence alignments, MAGs and annotations

https://doi.org/10.5281/zenodo.18410361

#### Input data and analyses script for figures and statistics

https://github.com/amankowski/placozoan_microbiome
